# Establishment and characterization of a brain cell line from bighead carp (*Hypophthalmichthys nobilis*)

**DOI:** 10.64898/2026.11.12.766164

**Authors:** Xiuying Guan, Yuan Lin, Jiameng Cui, Xinhang Chen, Dongming Li, Peng Jia

**Affiliations:** Fuzhou Medical College (former Fuzhou Medical College of Nanchang University), Fuzhou, Jiangxi 344000, China; Department of Respiratory and Critical Care Medicine, The First Affiliated Hospital of Nanchang University, Nanchang, Jiangxi 330006, China

**Keywords:** bighead carp, brain cell line, SVCV, viral susceptibility, innate immunity

## Abstract

A wide range of pathogens continues to threaten bighead carp (*Hypophthalmichthys nobilis*) aquaculture. Fish cell lines allow host–pathogen interactions to be dissected and candidate control measures to be tested under defined conditions. We established and characterized the first brain-derived cell line from bighead carp, designated HnB, which is only the second cell line reported for this species. HnB cells exhibited stable proliferation in Dulbecco’s Modified Eagle Medium (DMEM) containing 5–20% fetal bovine serum (FBS) at 28–32 °C and were serially passaged more than 50 times. The species origin of HnB cells was confirmed by PCR amplification and sequencing of the cytochrome b gene, and the sequence obtained showed 100% identity with previously published *H. nobilis cytochrome b* sequences. The modal chromosome number was 48, and the cells were free of mycoplasma contamination. Transfection with pEGFP-N3 demonstrated that HnB cells support exogenous gene expression. Quantitative real-time PCR (qRT-PCR) and viral titre assays showed that HnB cells were susceptible to spring viraemia of carp virus (SVCV). Moreover, both *interferon alpha* (*IFN-α)* and *interferon-stimulated gene 15* (*ISG15*) were significantly up-regulated in HnB cells following exposure to SVCV or poly(I:C), whereas LPS down-regulated the expression of both genes, indicating that HnB cells mount transcriptional antiviral responses to viral and viral-mimetic stimuli. The HnB line therefore supports SVCV replication and gives a species-matched system for studying antiviral (*IFN-α*/*ISG15*) responses in bighead carp.

## 1. Introduction

Bighead carp (*Hypophthalmichthys nobilis*), a member of the genus *Hypophthalmichthys* (family Cyprinidae), is widely distributed across China, from Hainan Island in the south to the Heilongjiang River Basin in the north, and is highly valued for its palatability and nutritional quality. It has long been recognized as one of the four major Chinese carps, alongside black carp, grass carp and silver carp, and remains one of the seven major freshwater aquaculture species in China (Wan et al. 2025). However, diseases continue to constrain bighead carp aquaculture, particularly as the industry becomes increasingly intensified and globalized. Although this species was once regarded as disease-resistant and highly productive, the emergence and spread of multiple pathogens have recently caused substantial losses. Spring viraemia of carp virus (SVCV), a notifiable pathogen listed by the World Organisation for Animal Health (WOAH; formerly OIE), is a major viral threat: the virus is characterized by rapid transmission, a short incubation period and high mortality, typically 30–40%, occasionally reaching 70% and approaching 100% in severe outbreaks (Shao and Zhao 2017). Therefore, elucidating the pathogenic mechanisms of SVCV and the antiviral responses of the host is essential for the management and prevention of this disease in bighead carp.

Fish cell lines provide valuable *in vitro* platforms for virology, immunology, toxicology and other fields (Kumar *et al*. 2023). For instance, a fin cell line from Salmo salar, a liver cell line from Acanthopagrus latus and a brain cell line from *Sparus aurata* have been used to investigate nervous necrosis virus, viral haemorrhagic septicaemia virus and SVCV, respectively (Jia et al. 2022a; Li et al. 2022; Ruiz-Palacios *et al*. 2020b). Kidney and brain cell lines from *Cromileptes altivelis* have shown susceptibility to *Vibrio harveyi* and *Edwardsiella tarda*, respectively, making them useful for studying bacterial cytotoxicity (Liu *et al*. 2021; Wei *et al*. 2022). Moreover, a brain cell line from *Lateolabrax japonicus* has been used to study innate immune responses to viral infection (Jia *et al*. 2022b; Whelan *et al*. 2020). As of January 2025, the Cellosaurus database catalogued 960 fish cell lines (Bairoch 2018; Cellosaurus 2025). Nevertheless, these lines represent only a small fraction of fish diversity: more than 37,000 valid fish species are currently recognized (Fricke *et al*. 2026). Species-specific models therefore remain scarce for most species, including economically important ones such as bighead carp.

To date, the only cell line reported from bighead carp is the snout-derived BHS line established in 1993, whose susceptibility to pathogens has not been characterized and whose modal chromosome number shifted from 48 to 71 between passages 15 and 25 (Shen et al. 1993). Given the economic importance of bighead carp and the reliance of functional studies on species-matched in vitro systems, a second, well-characterized cell line from this species represents a valuable resource.

Among candidate tissues, brain tissue is a particularly useful source. The brain is consistently invaded by SVCV and is recognized as one of its target tissues: SVCV RNA was detected most reliably in the brain among the organs examined by non-radioactive DNA probe hybridization (Oreshkova et al. 1999), and in situ hybridization confirmed viral localization in the brain of diseased fish (Wu et al. 2017). Accordingly, the brain is included among the sample tissues recommended by WOAH for SVCV diagnosis (WOAH 2023). Consistent with this tropism, brain-derived cell lines from other fish species, such as the gilthead seabream line SaB-1 and the mummichog line FuB-1, have proven permissive to SVCV and useful for studies of antiviral immunity (Ruiz-Palacios et al. 2020a, 2020b). Here we established a brain cell line from bighead carp (designated HnB) and evaluated its susceptibility to SVCV and its innate immune responses to SVCV, poly(I:C) and LPS. The line is offered as a well-characterized alternative to the snout-derived BHS line for work that requires cells of bighead carp origin.

## 2. Materials and Methods

### 2.1 Fish and virus

Healthy bighead carp (approximately 25 cm in length) were obtained from a commercial fish farm in Nancheng County, Fuzhou City, Jiangxi Province, China, and maintained at 25 °C. All experimental procedures involving fish were approved by the Ethics Committee of Nanchang University. Prior to dissection, fish were euthanised by immersion in an overdose of tricaine methanesulfonate (MS-222; Sigma, Germany). SVCV stocks were stored at −80 °C in our laboratory.

### 2.2 Primary cell culture and subculture

Primary cultures of bighead carp brain tissue were established using enzymatic digestion under strict aseptic conditions. The fish surface was sterilized with 75% ethanol, and brain tissue was carefully dissected using sterile scissors. The tissue was washed three times with phosphate-buffered saline (PBS), briefly immersed in 75% ethanol for 30 s and washed twice more with PBS. The brain tissue was then minced into approximately 1 mm³ fragments and digested with trypsin–EDTA (0.25%) for 20 min at room temperature. After centrifugation to remove the trypsin solution, the cells were resuspended in 4 mL of Dulbecco’s Modified Eagle Medium (DMEM) supplemented with 15% fetal bovine serum (FBS; CellMax, China) and transferred to a 25 cm² culture flask maintained at 28 °C. The culture medium was replaced every 2–3 days until the cells had adhered and formed a monolayer. Cells were then detached using trypsin and subcultured at a 1:2 ratio. This cell line was designated HnB.

### 2.3 Cell cryopreservation and recovery

Cryopreservation and recovery of HnB cells were performed as previously described (Li *et al*. 2019). For cryopreservation, HnB cells at approximately 90% confluence in a 25 cm² flask were trypsinized and resuspended in 1.8 mL of cryopreservation medium (10% dimethyl sulfoxide and 40% FBS in DMEM). The cell suspension was transferred to a 2 mL cryovial and sequentially cooled at 4 °C for 1 h, −20 °C for 2 h and −80 °C overnight, before being stored in a −150 °C ultra-low-temperature freezer (Haier, China).

For recovery, cryopreserved HnB cells were rapidly thawed in a 30 °C water bath and centrifuged at 1000 rpm for 5 min to remove the cryopreservation medium. The cell pellet was washed with DMEM, centrifuged again and resuspended in DMEM containing 15% FBS. Cell viability was assessed using trypan blue staining, and cells were seeded into a 25 cm² culture flask.

### 2.4 Species authentication

To confirm the species origin of HnB cells, a fragment of the *cytochrome b* (*cyt b*) gene was amplified by PCR. Genomic DNA was extracted from both bighead carp brain tissue and HnB cells (passage 10) using a Tissue DNA Kit (Vazyme, China). PCR amplification was performed using primers *cyt b*-F and *cyt b*-R (Table S1). The PCR products were cloned into the pMD-19T vector (TaKaRa, China) and sequenced. The obtained sequences were compared with those in the NCBI GenBank database. Species authentication and karyotyping were last performed at Fuzhou Medical College in May 2024.

### 2.5 Mycoplasma detection

Mycoplasma contamination was assessed using the Myco-Lumi™ Luminescent Mycoplasma Detection Kit (C0297; Beyotime, China) according to the manufacturer’s instructions. This luminescent assay is based on the activity of mycoplasma-specific enzymes that convert ADP to ATP, which is quantified by a luciferase-dependent reaction. HnB cells at passage 45 were cultured for 5 days, and 200 µL of culture supernatant was clarified by centrifugation at 200 × g for 5 min to remove floating cells and debris. Fifty microlitres of the supernatant, the positive control supplied with the kit and fresh complete culture medium (negative control) were each mixed with 50 µL of detection reagent A in a white 96-well plate, and luminescence was recorded after 5 min at room temperature (reading A). Detection reagent B (50 µL) was then added, and luminescence was recorded after a further 10 min (reading B). The B/A ratio was calculated, and ratios below 0.9 were considered negative for mycoplasma contamination. Mycoplasma testing was last performed at Fuzhou Medical College in November 2026.

### 2.6 Chromosome analysis

Chromosome analysis of HnB cells at passage 40 was performed according to an adapted protocol (Li *et al*. 2022). Cells at 60–80% confluence were treated with colchicine (Sigma, Germany) for 6 h, then trypsinized and incubated in a hypotonic solution (0.075 M KCl) at 37 °C for 35 min. Cells were fixed twice with fresh Carnoy’s fixative (10 min each), dropped onto pre-chilled slides, air-dried and stained with 10% Giemsa (Sigma, Germany) for 10 min. At least 50 metaphase spreads were examined under a light microscope (Olympus, China).

### 2.7 Growth curves

Growth characteristics of HnB cells at passage 30 were evaluated under different temperatures and FBS concentrations. Cells were seeded into 24-well plates and cultured in DMEM with 15% FBS at 24, 28 or 32 °C. To assess the effect of FBS concentration, cells were cultured at 28 °C in DMEM containing 5%, 10%, 15% or 20% FBS. Cell numbers were counted daily for 7 days using a hemocytometer. All experiments were performed in triplicate.

### 2.8 Transfection

HnB cells (passage 45) were seeded into 12-well plates and cultured at 28 °C until reaching approximately 70% confluence. A total of 1 µg of the pEGFP-N3 plasmid was transfected into the cells using Lipofectamine™ 8000 (Beyotime, China) according to the manufacturer’s instructions. Green fluorescence signals were observed under an Olympus IX73 microscope 48 h post-transfection. Transfection efficiency was estimated by counting the proportion of fluorescent cells in 20 randomly selected fields.

### 2.9 Virus susceptibility and viral replication

HnB cells were seeded into 6-well plates at a density of 1 × 10^5^ cells per well and incubated at 28 °C for 24 h. Cells were then infected with SVCV at a multiplicity of infection (MOI) of 1, while control cells were treated with PBS. Cytopathic effects (CPE) were monitored daily. Cell samples were collected at 24 and 48 h post-infection (hpi) for RT-PCR and qRT-PCR analysis to evaluate viral replication, and cell supernatants were used for viral titre assay. Fragments of the SVCV *genes encoding the glycoprotein (G), RNA polymerase (L), matrix protein (M), nucleoprotein (N) and phosphoprotein (P)* were amplified with primers listed in Table S1.

### 2.10 Transcription of immune genes in HnB cells treated with SVCV, poly(I:C) and LPS

HnB cells were seeded into 6-well plates at a density of 1 × 10^5^ cells per well and incubated at 28 °C for 24 h. Cells were then stimulated with SVCV (MOI of 1), poly(I:C) (5 µg/mL, InvivoGen) or LPS (100 ng/mL, Beyotime). Total RNA was extracted at 24 and 48 h post-stimulation, and qRT-PCR was performed to assess the mRNA levels of interferon alpha (*IFN-α*) and interferon-stimulated gene 15 (*ISG15*).

### 2.11 RT-PCR

Total RNA was isolated from SVCV-infected and control HnB cells using TRIzol reagent (Invitrogen, USA) and reverse-transcribed into cDNA using the PrimeScript™ First-Strand cDNA Synthesis Kit (TaKaRa, China) according to the manufacturer’s instructions. PCR conditions were as follows: initial denaturation at 94 °C for 5 min; 30 cycles of 94 °C for 30 s, 55 °C for 30 s and 72 °C for 30 s, followed by a final extension at 72 °C for 10 min. PCR products were cloned into the pMD-19T vector and confirmed by sequencing.

### 2.12 qRT-PCR

qRT-PCR was performed using a BIO-RAD CFX Connect system. The thermal cycling conditions included an initial denaturation at 95 °C for 2 min, followed by 40 cycles of 95 °C for 15 s and 60 °C for 30 s. The *β-actin* gene was used as the internal reference gene. Relative gene expression levels were calculated using the 2^−ΔΔCt^ method. All samples were analyzed in triplicate.

### 2.13 Viral titre determination (TCID50 assay)

The viral titre was quantified using the 50% tissue culture infectious dose (TCID50) assay as previously described (Pan *et al*. 2024). Briefly, HnB cells were seeded into 96-well plates and grown to 80–90% confluence. The collected supernatants were serially diluted 10-fold (from 10^−1^ to 10^−8^) in DMEM. Aliquots (100 μL) of each dilution were added to the 96-well plates, with eight replicates per dilution. After 1 h of adsorption at 28 °C, the virus inoculum was replaced with fresh DMEM containing 2% FBS, and the cells were further incubated at 28 °C. CPE were monitored daily for 4–7 days. The viral titre was calculated using the Reed-Muench method and expressed as log_10_ TCID50/mL.

### 2.14 Statistical analysis

Data are presented as the mean ± SD from three independent experiments. Statistical analyses were performed using SPSS version 20. An unpaired Student’s t-test was used for two-group comparisons, and one-way ANOVA followed by Tukey’s post-hoc test was applied for multiple-group comparisons. *P* < 0.05 was considered statistically significant.

## 3. Results

### 3.1 Primary cell culture and subculture of HnB cells

The HnB cell line derived from the brain of healthy bighead carp was established by enzymatic digestion and maintained in DMEM supplemented with 15% FBS. By day 2, a small number of cells had adhered to the bottom of the culture flask, and the cells reached confluence after approximately 8 days. Subsequently, the cells were trypsinized and passaged at a 1:2 ratio every 2–3 days. Fibroblast-like, neuron-like and epithelial-like cells were identified in the primary cultures, with neuron-like cells growing above the epithelial-like cells (Figure 1A). In the early passages, epithelial-like, neuron-like and fibroblast-like cells were all observed (Figures 1B and 1C). During serial passaging, the neuron-like and epithelial-like cells gradually disappeared, and by passage 15 only fibroblast-like cells remained; the cell line has since been maintained for more than 50 passages (Figure 1D).

**FIGURE 1.**
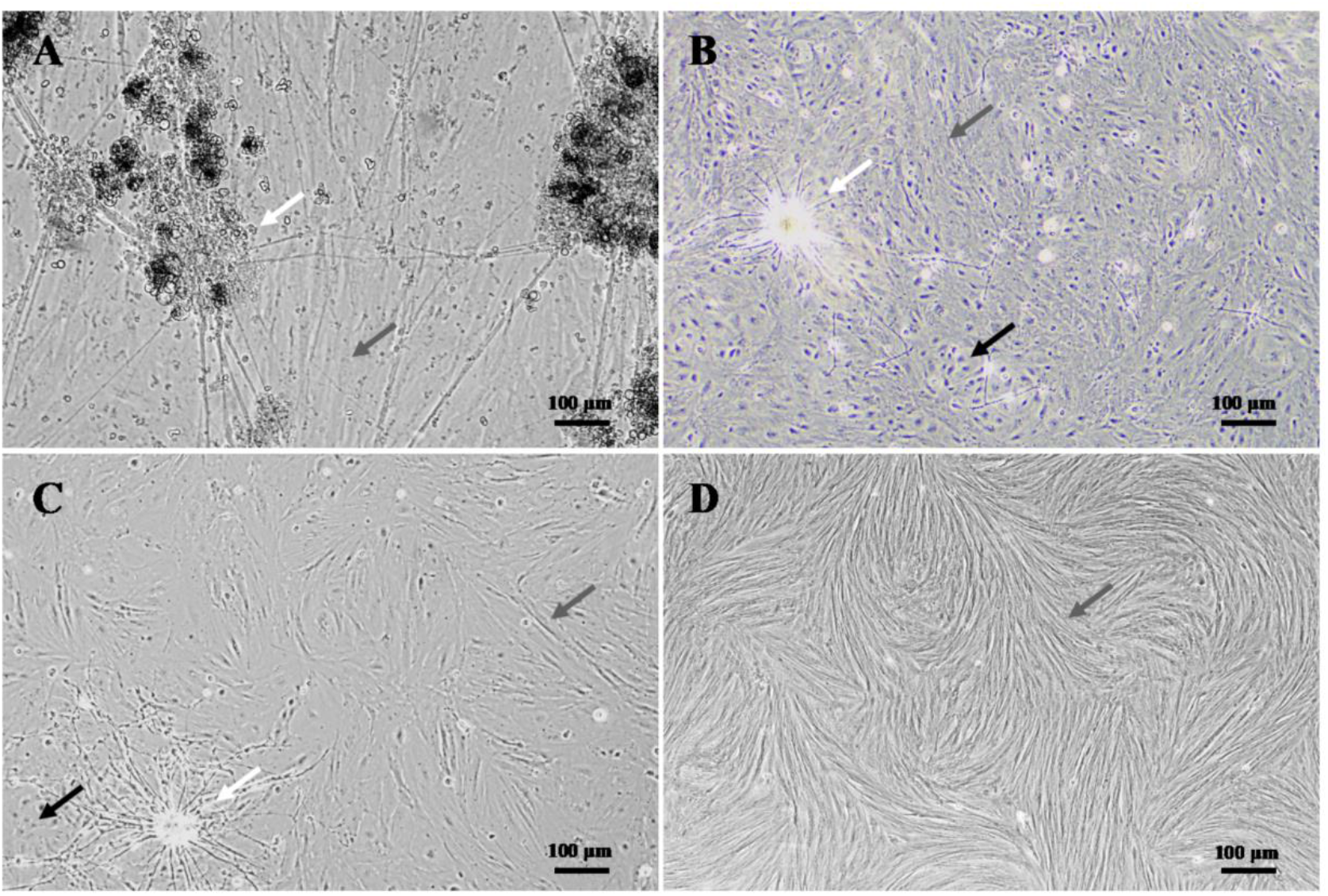
Primary culture and subculture of bighead carp brain cells. (A) HnB cells in primary culture on day 8; (B) HnB cells at passage 2; (C) HnB cells at passage 8; (D) HnB cells at passage 50. Black arrows indicate epithelial-like cells, white arrows indicate neuron-like cells, and gray arrows indicate fibroblast-like cells. Scale bar, 100 μm.

### 3.2 Species authentication and karyotyping

A 393-bp fragment of the conserved region of the bighead carp *cyt b* gene was amplified from genomic DNA extracted from both bighead carp brain tissue and HnB cells. The PCR products were analyzed by agarose gel electrophoresis (Figure 2A), revealing two bands of the expected size. The bands were excised, ligated into pMD-19T and sequenced. The obtained sequence showed 100% identity with the bighead carp *cyt b* sequence deposited in GenBank (accession no. MW862069.1), confirming the bighead carp origin of HnB cells. Chromosome counting of 50 metaphase spreads from HnB cells revealed a diploid chromosome number of 48 (2n = 48, n = 24) (Figures 2B and 2C). In addition, mycoplasma testing of HnB culture supernatant yielded a B/A ratio of 0.87, below the 0.9 threshold, whereas the positive control yielded 3.20, indicating that HnB cells were free of mycoplasma contamination.

**FIGURE 2.**
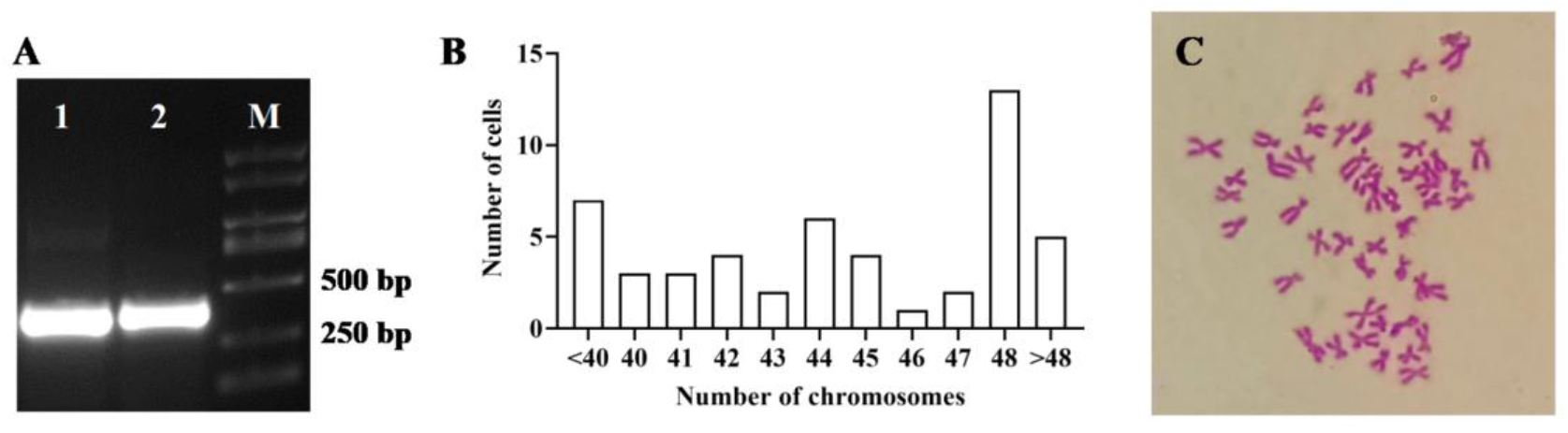
Species authentication and karyotyping of HnB cells. (A) Agarose gel electrophoresis of the *cyt b* gene PCR products (393 bp). M, DL2000 DNA marker; Lane 1, bighead carp brain tissue; Lane 2, HnB cells. (B) Chromosome number distribution of HnB cells. (C) Giemsa-stained metaphase chromosomes of HnB cells.

### 3.3 Cell growth characteristics

The effects of temperature and FBS concentration on the growth of HnB cells at passage 30 were investigated. Cell growth increased in a concentration-dependent manner over the range of 5–20% FBS, and proliferation remained relatively high even at 5% FBS (Figure 3A). With respect to temperature, HnB cells exhibited consistently high proliferation rates at both 28 and 32 °C, whereas proliferation nearly ceased at 24 °C (Figure 3B).

**FIGURE 3.**
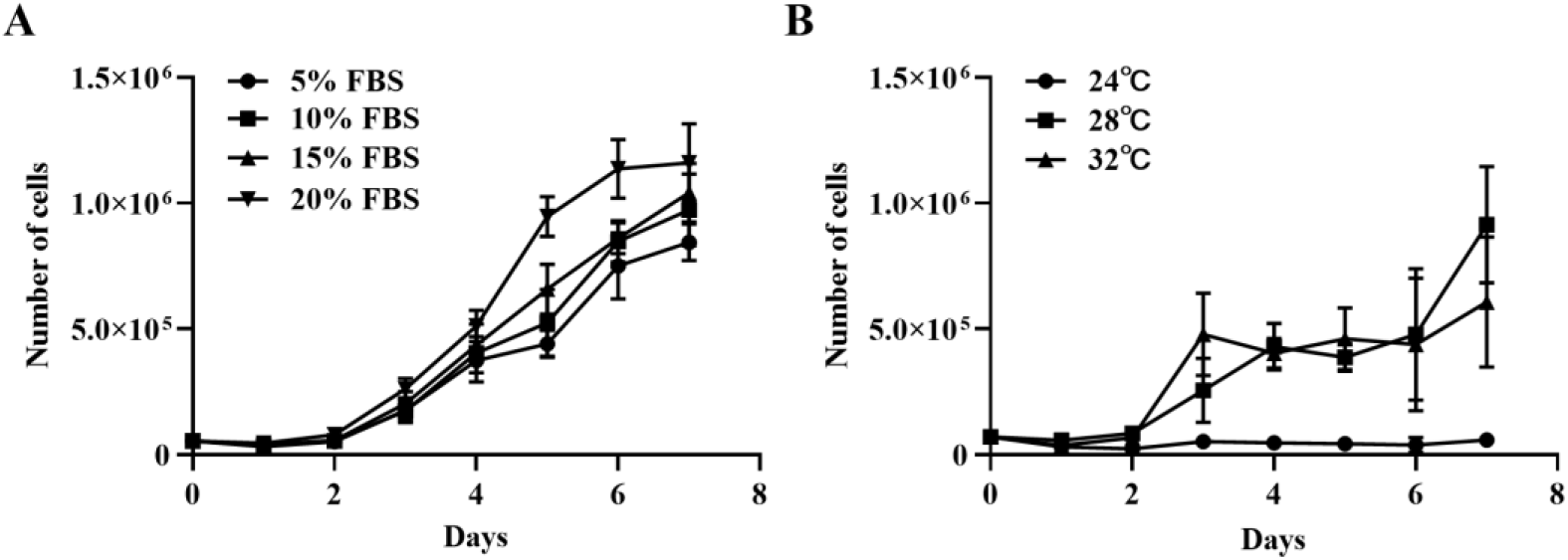
Growth curves of the HnB cell line. (A) Growth curves of HnB cells cultured at different FBS concentrations. (B) Growth curves of HnB cells cultured at different temperatures.

### 3.4 Transfection efficiency

To evaluate the suitability of HnB cells for exogenous gene manipulation, transfection with the pEGFP-N3 plasmid was performed. Green fluorescent signals indicative of successful transfection were observed 48 h post-transfection, and the transfection efficiency was estimated to be approximately 5% based on cell counting (Figure 4).

**FIGURE 4.**
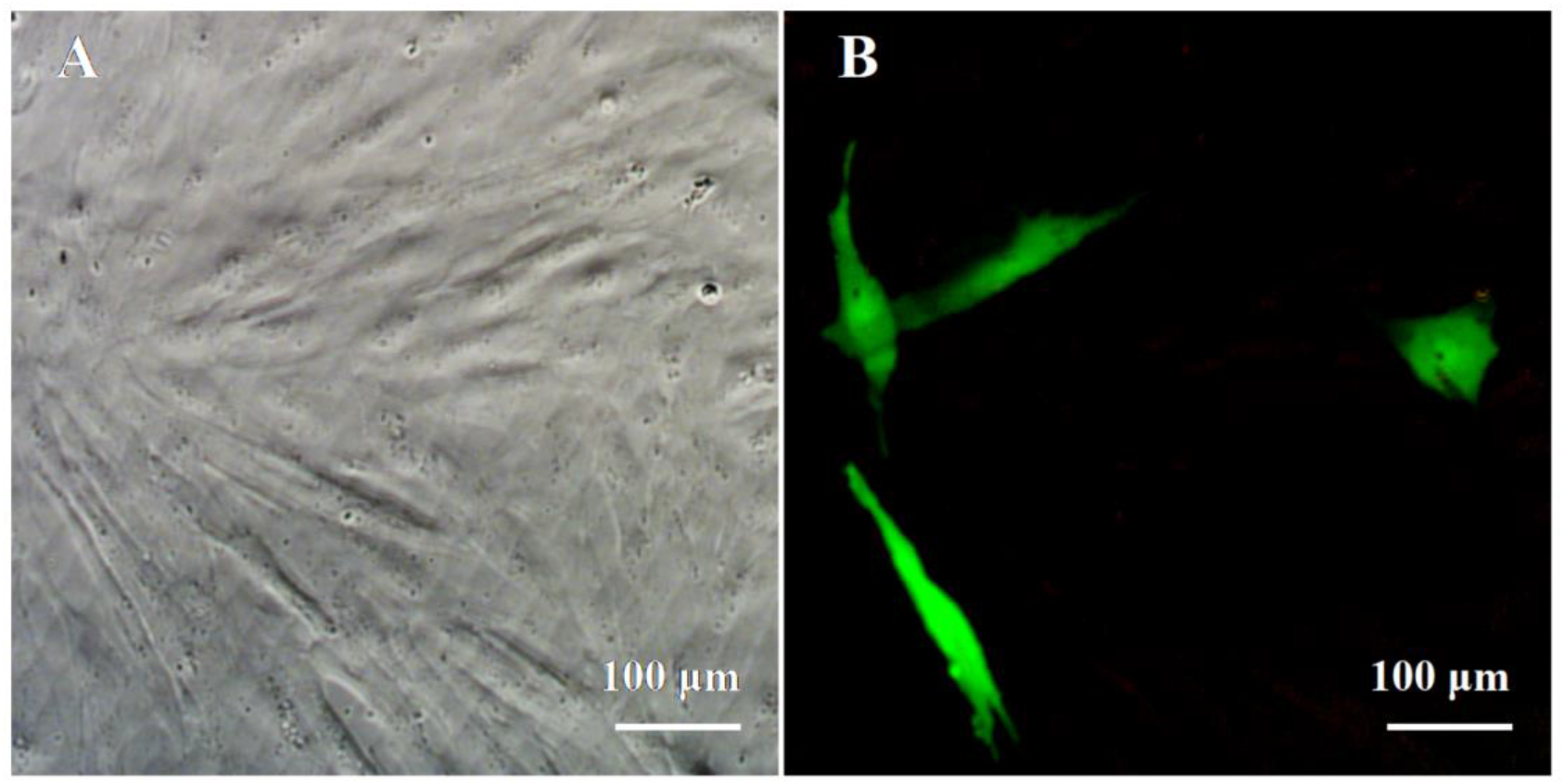
**Transfection of HnB cells with pEGFP-N3**. HnB cells were observed under bright-field (A) and fluorescence (B) microscopy 48 h post-transfection.

### 3.5 Viral susceptibility and viral replication

The susceptibility of HnB cells to SVCV was assessed. Compared with control cells (Figures 5A and 5B), SVCV-infected cells displayed characteristic cytopathic effects at 24 and 48 hpi (Figures 5C and 5D), including vacuolation, shrinkage, rounding and detachment from the culture surface. In addition, a partial sequence of the SVCV *G* gene (396 bp) was amplified from SVCV-infected HnB cells (Figure 5E). The replication of SVCV in HnB cells was determined by qRT-PCR; transcript levels of the SVCV *G*, *L*, *M*, *N* and *P* genes increased significantly from 24 to 48 hpi in infected cells (Figure 5F).

**FIGURE 5.**
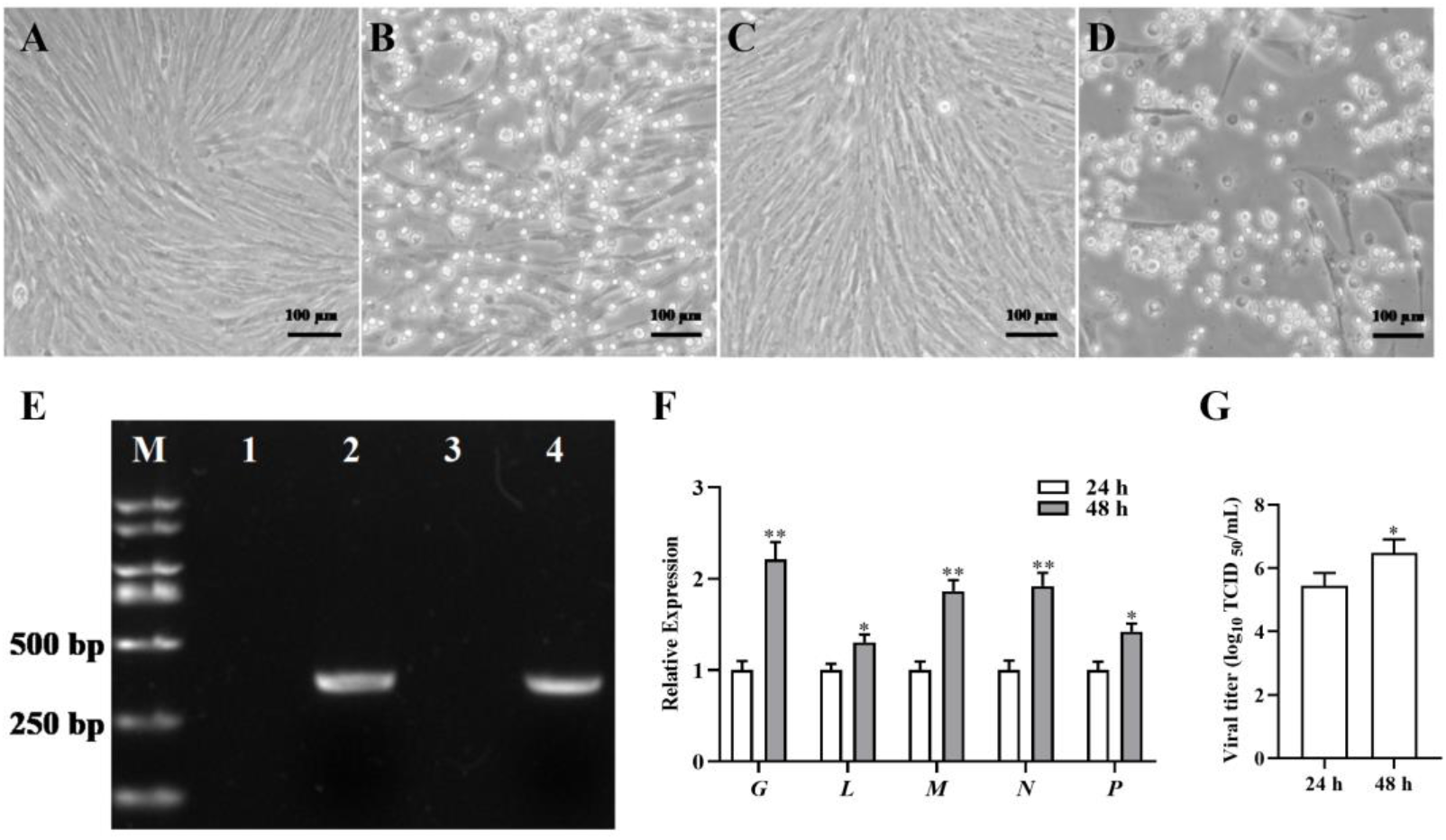
Susceptibility of HnB cells to SVCV. (A, B) Mock-infected HnB cells at 24 and 48 hpi. (C, D) Cytopathic effects (CPE) in SVCV-infected HnB cells at 24 and 48 hpi. (E) Agarose gel electrophoresis of PCR products from SVCV-infected HnB cells using SVCV *G* gene-specific primers. M, DL2000 DNA marker; Lane 1, mock-infected HnB cells at 24 hpi; Lane 2, SVCV-infected HnB cells at 24 hpi; Lane 3, mock-infected HnB cells at 48 hpi; Lane 4, SVCV-infected HnB cells at 48 hpi. (F) qRT-PCR analysis of SVCV gene (*G*, *L*, *M*, *N* and *P*) expression in SVCV-infected HnB cells at 24 and 48 hpi. *, *P <* 0.05; **, *P <* 0.01. (G) Viral titres in the supernatants of SVCV-infected HnB cells at 24 and 48 hpi, determined by TCID50 assay. Data are presented as the mean ± SD of three independent experiments (n = 3). *, *P <* 0.05.

To further quantify the replication dynamics of SVCV in HnB cells, the viral titres in the culture supernatants were determined at 24 and 48 hpi using the TCID50 assay. As shown in Figure 5G, the viral titre increased significantly from 5.44 ± 0.40 log10 TCID50/mL at 24 hpi to 6.48 ± 0.42 log10 TCID50/mL at 48 hpi (*P* < 0.05). The titre thus rose by about one log10 unit between 24 and 48 hpi, in line with the progression of CPE and the accumulation of viral transcripts.

### 3.6 Expression of immune genes in HnB cells induced by SVCV, poly(I:C) and LPS

To examine the response of HnB cells to pathogen stimulation, the expression levels of *IFN-α* and *ISG15* were assessed in HnB cells treated with SVCV, poly(I:C) or LPS. Compared with the control group, the transcript levels of *IFN-α* and *ISG15* were significantly up-regulated in HnB cells at 24 and 48 h after stimulation with SVCV (Figures 6A and 6B) or poly(I:C) (Figures 6C and 6D). In contrast, down-regulation of these transcripts was observed in cells stimulated with LPS (Figures 6E and 6F).

**FIGURE 6.**
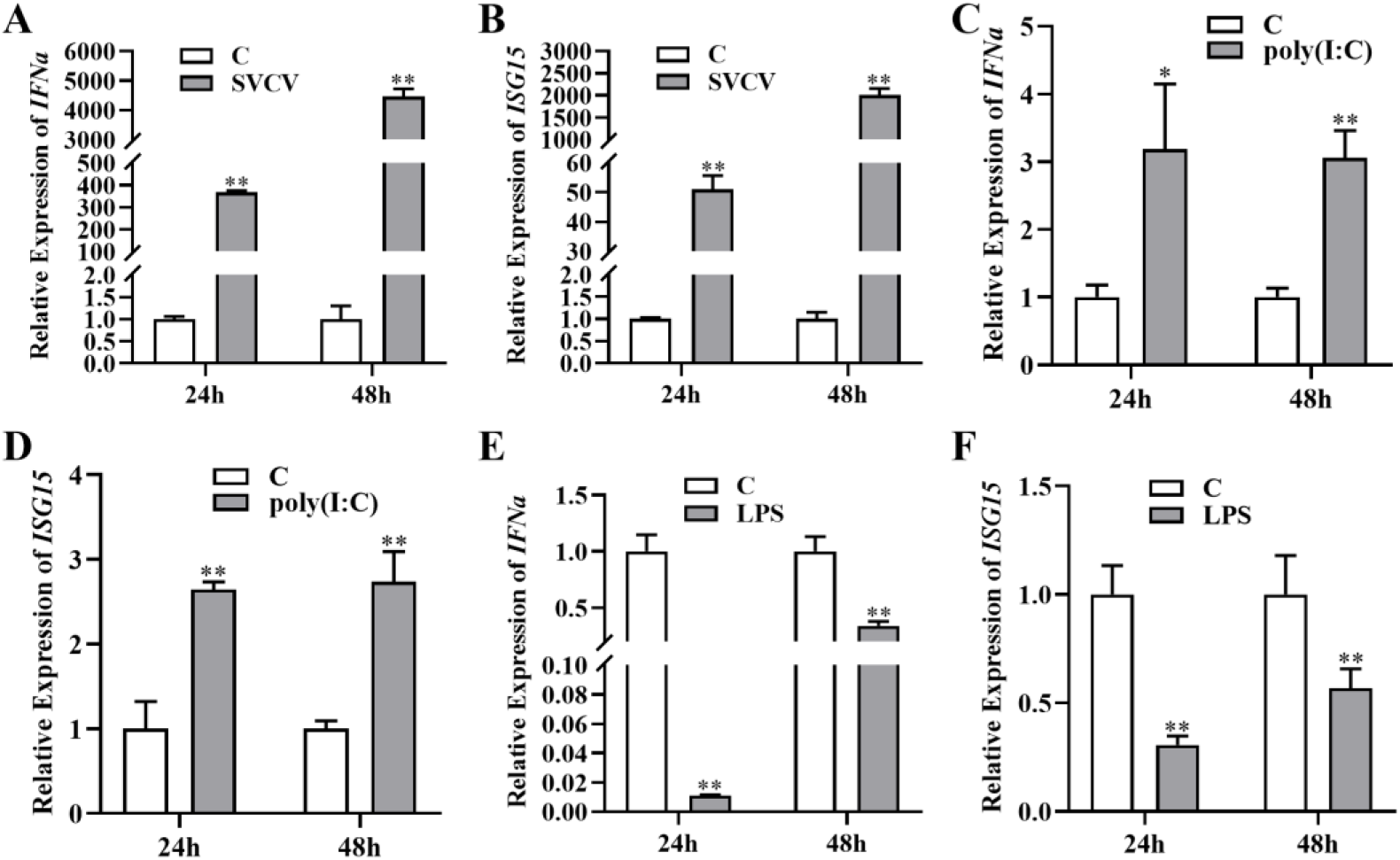
Expression of immune genes in HnB cells after SVCV, poly(I:C) or LPS stimulation. (A, B) Transcript levels of *IFN-α* and *ISG15* in HnB cells at 24 and 48 h after SVCV infection. (C, D) Transcript levels of *IFN-α* and *ISG15* in HnB cells at 24 and 48 h after poly(I:C) stimulation. (E, F) Transcript levels of *IFN-α* and *ISG15* in HnB cells at 24 and 48 h after LPS stimulation. Data are presented as the mean ± SD of three independent experiments performed in triplicate. *, *P* < 0.05; **, *P* < 0.01.

## 4. Discussion

In this study, a cell line designated HnB was established from the brain tissue of bighead carp (*Hypophthalmichthys nobilis*) by enzymatic digestion, a method reported to yield cells more rapidly than the tissue block culture method (Deng et al. 2023). HnB cells have been maintained for more than 50 passages in DMEM, whereas many fish brain cell lines are cultured in Leibovitz’s L-15 medium (Liao et al. 2023; Liu et al. 2021; Ruiz-Palacios et al. 2020b). Although DMEM has been widely used for the propagation of mammalian glial and neuronal cells (Loui et al. 2025), the present results show that it also supports the long-term cultivation of fish brain cells. As in a previous report (Liao et al. 2023), neuron-like and epithelial-like cells disappeared during serial passaging, and by passage 15 only fibroblast-like cells remained, presumably because these cell types proliferated slowly or not at all and were therefore lost during passaging. Accordingly, HnB is best suited to virological and antiviral-immunity studies rather than to studies requiring neuronal properties.

PCR amplification of the cyt b gene confirmed the bighead carp origin of HnB cells. Although the modal chromosome number of HnB cells at passage 40 was 48 (2n = 48, n = 24), consistent with that reported for bighead carp (Jian et al. 2020), a substantial proportion of metaphases (37/50, 74%) exhibited numerical variations ranging from 40 to 48 (Figure 2B). This extensive chromosomal heterogeneity is a common hallmark of long-term in vitro culture and indicates a degree of chromosomal instability within the cell line. By comparison, the previously reported bighead carp snout cell line (BHS) maintained a modal number of 48 at passage 15 but shifted to 71 at passage 25 (Shen et al. 1993). The HnB line retained 48 as the modal number at passage 40 despite this variation, indicating that it is relatively more stable than BHS.

To determine the optimal growth conditions, we assessed the effects of FBS concentration and temperature on the proliferation of HnB cells. Cell proliferation increased with FBS concentration over the range of 5–20% but remained appreciable even at 5% FBS, a trend consistent with observations in the *Epinephelus coioides* spleen cell line and the *Pampus argenteus* liver cell line (Huang *et al*. 2018; Jiang *et al*. 2023). HnB cells maintained high proliferation rates at both 28 and 32 °C, whereas growth nearly ceased at 24 °C. The BHS cell line, derived from the snout of bighead carp, showed an optimal temperature range of 25–30 °C (Shen *et al*. 1993); the two ranges overlap between 28 and 30 °C, and the apparent difference may reflect differences in tissue origin.

Transfection with the pEGFP-N3 plasmid was achieved in HnB cells using a liposome-based system, with an estimated efficiency of approximately 5%, indicating that HnB cells support liposome-mediated gene delivery.

Fish cell lines are useful tools for *in vitro* virological research (Deng *et al*. 2024; Dinakaran *et al*. 2024; Ghermezian *et al*. 2024). Here, we evaluated the susceptibility of HnB cells to SVCV, a negative-sense single-stranded RNA virus belonging to the genus *Sprivivirus* (family Rhabdoviridae) that causes high mortality in several cyprinid species, including bighead carp (Yin *et al*. 2024). Several cell lines from other fish species have been reported to be susceptible to SVCV, including a zebrafish caudal fin cell line (Yin *et al*. 2024), a gilthead seabream brain cell line (Ruiz-Palacios *et al*. 2020b) and a cell line derived from mummichog (*Fundulus heteroclitus*) brain (Ruiz-Palacios *et al*. 2020a). However, a systematic search of the Cellosaurus and Web of Science databases revealed no brain cell line derived from *Hypophthalmichthys nobilis*. The HnB cell line supported SVCV replication, as evidenced by the development of CPE, the detection of viral gene transcripts and an increase in viral titre to 6.48 log10 TCID50/mL at 48 hpi, a magnitude comparable to titres reported in other susceptible fish cell lines (Ruiz-Palacios et al. 2020a, 2020b). Rhabdoviruses of different genera share tetraspanin CD9 as a conserved entry receptor (Zhang *et al*. 2026), and the permissiveness of HnB cells observed here is consistent with this conserved entry pathway. Although widely used SVCV-permissive lines such as Epithelioma papulosum cyprini (EPC) cells are highly efficient for virus propagation, they are derived from Cyprinus carpio. HnB allows SVCV–host interactions and innate immune responses to be examined in cells of the same species as the host, rather than in a heterologous cyprinid line. The brain tropism of SVCV in vivo (Fijan et al. 1971; Oreshkova et al. 1999; Wu et al. 2017) supports the use of a brain-derived line as a substrate for examining the interaction of SVCV with brain-derived cells.

Cell lines are also widely used to study immune mechanisms in fish. Compared with animal models, fish cell lines provide a more convenient and efficient *in vitro* platform for gene function studies. Although many fish cell lines have been established and applied to this purpose (Lu *et al*. 2023; Yi *et al*. 2025; Zhang *et al*. 2022), interspecies differences make it preferable to use a cell line derived from bighead carp to investigate the functions of its immune genes. The innate immune response is the first line of defence against pathogens in fish. Upon pathogen entry, pattern recognition receptors detect pathogen-associated molecular patterns (PAMPs) and trigger the expression of interferons (IFNs) through signaling pathways activated by receptors such as the RIG-I-like receptors (RLRs) and Toll-like receptors (TLRs); IFNs in turn induce the expression of interferon-stimulated genes (ISGs), including ISG15, Mx-1 and viperin, which act to eliminate invading pathogens (Ortega-Villaizan et al. 2022). We examined the responses of HnB cells to SVCV, poly(I:C) and LPS by measuring the expression of *IFN-α* and *ISG15*. A previous study showed that SVCV infection induces the expression of *IFN-α* and *ISG15* in *EPC* cells (Qiu *et al*. 2022). Consistent with that finding, both *IFN-α* and *ISG15* were significantly up-regulated in SVCV-infected HnB cells at 24 and 48 h post-infection. Stimulation with poly(I:C), a synthetic analogue of double-stranded RNA commonly used to mimic viral infection, also significantly elevated the expression of both genes relative to the control, showing that HnB cells can mount an antiviral transcriptional response.

In contrast, the transcript levels of *IFN-α* and *ISG15* were down-regulated in LPS-stimulated HnB cells. LPS is a major component of the outer membrane of Gram-negative bacteria and is commonly used to mimic bacterial infection. The induction of type I interferons during bacterial infection may differ from that during viral infection. For example, LPS from Salmonella Typhimurium suppresses macrophage IFN-β signaling in a TLR4-dependent manner (Ando et al. 2015; Shuster et al. 2024), and in sea perch (*Lateolabrax japonicus*), the expression of MDA5, an upstream signaling molecule in the IFN pathway, is also suppressed by LPS (Jia et al. 2016). Bacterial and viral stimuli therefore act on HnB cells through different routes, although the mechanism behind this difference remains to be defined.

Two points limit the present characterization. Karyotype and growth were each assessed at a single passage (P40 and P30, respectively), so chromosomal drift during prolonged culture remains untracked and periodic checks would be needed before the line is used for cytogenetic work. Transfection with Lipofectamine 8000 reached only about 5%, adequate for transient reporter assays but too low for stable-line work; altering the liposome:DNA ratio or switching to electroporation may improve it. HnB nonetheless provides a workable in vitro system for virus isolation, host–pathogen interaction studies and functional analysis of immune genes in bighead carp.

## 5. Conclusion

The HnB line described here is the second cell line from bighead carp and the first derived from its brain. HnB cells supported SVCV replication, and the transcripts of IFN-α and ISG15 rose after stimulation with SVCV and poly(I:C), whereas LPS suppressed both genes. As a karyotyped, mycoplasma-free line of bighead carp origin, HnB provides a platform for virus isolation and for dissecting IFN-α/ISG15 responses in this species.

## Author Contributions

X.G., Y.L., J.C. and X.C. performed the experiments and analyzed the data. X.G. wrote the first draft of the manuscript. D.L. and P.J. acquired the funding, conceived and designed the study, and participated in the writing and revision of the manuscript. All authors read and approved the final manuscript.

## Data Availability Statement

The original data generated and analyzed in this study are available from the corresponding author upon reasonable request.

## Compliance with Ethical Standards

Conflict of Interest Author Xiuying Guan declares no conflict of interest. Author Yuan Lin declares no conflict of interest. Author Jiameng Cui declares no conflict of interest. Author Xinhang Chen declares no conflict of interest. Author Dongming Li declares no conflict of interest. Author Peng Jia declares no conflict of interest.

## Supporting information

Supplemental Table 1. Primers used in this study

