## Supplemental Table 1. Primers used in this study for "Establishment and characterization of a brain cell line from bighead carp (*Hypophthalmichthys nobilis*)"

**Table S1** Sequences of primers used in this paper

| Name | Sequence（5'-3'） | Comment |
|---|---|---|
| *Cyt b*-F | AGCCTACGAAAAACCCACCC | Partial sequence of *Cyt B* |
| *Cyt b*-R | AAGGACGTAGCCAACGAAGG | |
| SVCV-*G*-F | TGAAGAYTGTGTCAATCAAGTC | Partial sequence of SVCV *G* |
| SVCV-*G*-R | GCGARTGCAGAGAAAAAGTG | |
| *β-actin*-qF | TCGTCCACCGCAAATGCTTCTA | QPCR of *β-actin* |
| *β-actin*-qR | CCGTCACCTTCACCGTTCCAGT | |
| SVCV-*N*-qF | TGAGTGCTGAGGACGAT | QPCR of SVCV *N* |
| SVCV-*N*-qR | TTTGTGAGTTGCCGTTA | |
| SVCV-*P*-qF | TTGGACCTGGGATAGTGA | QPCR of SVCV *P* |
| SVCV-*P*-qR | CTTGCTTGGTTTGTGGG | |
| SVCV-*M*-qF | TACTCCTCCCACTTACGA | QPCR of SVCV *M* |
| SVCV-*M*-qR | CAAGAGTCCGAGAAGGTC | |
| SVCV-*G*-qF | CGACCTGGATTAGACTTG | QPCR of SVCV *G* |
| SVCV-*G*-qR | AATGTTCCGTTTCTCACT | |
| SVCV-*L*-qF | GCCCACTTTGCATCCAGTCC | QPCR of SVCV *L* |
| SVCV-*L*-qR | GAGATGCCACAGACTCCTCC | |
| *IFNα*-F | CCAGTGGGGTCCAAAAGAAT | QPCR of *IFNα* |
| *IFNα*-R | GCAGCTTGAAGTTCTTGACCAG | |
| *ISG15*-F | TCGCTGGATGTGAGAGGTAA | QPCR of *ISG15* |
| *ISG15*-R | TCTCGTTAATGACGAACACCTGAA | |
